# Melatonin Targets MoIcl1 and Works Synergistically with Fungicide Isoprothiolane in Rice Blast Control

**DOI:** 10.1101/2023.07.01.547317

**Authors:** Ruiqing Bi, Renjian Li, Zhenyi Xu, Huanyu Cai, Juan Zhao, Yaru Zhou, Bangting Wu, Peng Sun, Wei Yang, Lu Zheng, Xiao-Lin Chen, Chao-Xi Luo, Huailong Teng, Qiang Li, Guotian Li

## Abstract

Melatonin-a natural harmless molecule-displays versatile roles in human health and crop disease control such as for rice blast. Rice blast, caused by the filamentous fungus *Magnaporthe oryzae*, is one devastating disease of rice. Application of fungicides is one of the major measures in the control of various crop diseases. However, fungicide resistance in the pathogen and relevant environmental pollution are becoming serious problems. By screening for possible synergistic combinations, here, we discovered an eco-friendly combination for rice blast control, melatonin and the fungicide isoprothiolane. These compounds together exhibited significant synergistic inhibitory effects on vegetative growth, conidial germination, appressorium formation, penetration, and plant infection by *M. oryzae*. The combination of melatonin and isoprothiolane reduced the effective concentration of isoprothiolane by over 10-fold as well as residual levels of isoprothiolane. Transcriptomics and lipidomics revealed that melatonin and isoprothiolane synergistically interfered with lipid metabolism by regulating many common targets, including the predicted isocitrate lyase-encoding gene *MoICL1*. Furthermore, we show that melatonin and isoprothiolane interact with MoIcl1 using different techniques. This study demonstrates that melatonin and isoprothiolane function synergistically and can be used to reduce the dosage and residual level of isoprothiolane, potentially contributing to the environment-friendly and sustainable control of crop diseases.

## 1 INTRODUCTION

Melatonin (N-acetyl-5-methoxytryptamine), a naturally occurring tryptophan-derived indoleamine hormone, regulates various biological processes.^1^ Additionally, melatonin can inhibit the growth of pathogens and cancer cells.^2–5^ Exogenous application of melatonin confers plants resistance to cassava bacterial wilt, grape anthracnose, tobacco mosaic virus, watermelon powdery mildew, and so on.^6–10^ Regarding rice diseases, melatonin effectively inhibits bacterial leaf blight, rice stripe disease and rice blast.^11–13^

Rice (*Oryza sativa*) is a major staple crop in the world, providing food for more than half of the world’s population.^14^ However, the quantity and quality of rice are seriously threatened by various fungal diseases. Among them, rice blast caused by the filamentous fungus *Magnaporthe oryzae* alone, reduces global production of rice with the amount sufficient to feed over 60 million people.^15^ *M. oryzae* infects plants mainly through its asexual spores, conidia.^16^ Germinated conidia develop infection-specific structures called appressoria that play a vital role in plant infection. Fatty acids, glycogen, glycerol, and triacylglycerol are involved in the accumulation of enormous turgor pressure in the appressorium, which directly penetrates into the plant cell.^17–19^ Accordingly, inactivation of the enzymes involved in the metabolism of these important fatty acid and lipid species blocks critical pathways and suppresses plant infection.^18,20^

Fungicide application is often necessary for effective disease management.^21^ Fungicides including carbendazim, isoprothiolane, thiophanate methyl, and tricyclazole, are commonly used for rice blast control. Isoprothiolane inhibits the synthesis of phospholipid phosphatidylcholine,^22,23^ an important component of fungal cell membranes. Tricyclazole inhibits melanin biosynthesis of *M. oryzae*, which is crucial for the formation and penetration of appressorium.^24^ Thiophanate methyl inhibits hyphal growth and causes abnormal germ tube formation, suppressing infection.^25^ Carbendazim mainly interferes with mitotic spindles formation during cell division, inhibiting mycelial growth.^26^ Two million tons of fungicides are applied worldwide annually.^27^ Overuse of fungicides leads to environmental degradation, threatening human and animal health as well as biodiversity.^28,29^ Therefore, novel methods of fungicide application must be explored to effectively control crop diseases including rice blast, in an environment-friendly manner.

Fungicide combinations could be a powerful method to expand the antifungal spectrum of fungicides and reduce the dosage of fungicides, particularly if the combinations are synergistic, thereby reducing the cost, potential toxicity, and negative impacts of fungicides on the environment. Drug synergy is the result of the simultaneous functions of two modes of action or components.^30,31^ Many fungicide combinations have been successfully applied. For example, combined application of a biocontrol agent TA-1 and fungicide fluopimomide synergistically improves the control of gray mold in tomato.^32^ The synergistic effects of melatonin and various drugs have been reported in cancer treatment and disease management.^33–38^ Melatonin combined with vincristine or ifosfamide has a synergistic effect in inhibiting Ewing sarcoma;^33^ melatonin and selenium synergistically inhibit the growth of the fungal pathogen *Botrytis cinerea.*^35^

In this study, we demonstrated that the synergy of melatonin and isoprothiolane inhibited the growth of *M. oryzae*. Melatonin and isoprothiolane not only had synergistic inhibitory effects on development but also plant infection by *M. oryzae*. Transcriptomics and lipidomics revealed that melatonin and isoprothiolane synergistically interfered with lipid metabolism by suppressing the activity of many common targets. Furthermore, we predicted that both melatonin and isoprothiolane target the isocitrate lyase-encoding protein MoIcl1, and we confirmed the interactions using surface plasmon resonance (SPR) and nano differential scanning fluorimetry (Nano DSF) assays. Overall, this study shows that synergistic action of melatonin and isoprothiolane controls rice blast, potentially reducing the amount and residual levels of fungicides, and increasing the levels of melatonin in rice.

## 2 MATERIALS AND METHODS

### 2.1 Strains and culture conditions

*M. oryzae* strains were routinely maintained at 28 ℃ on oatmeal agar medium (OMA) or complete medium (CM).^39^ The rice (*Oryza sativa*) variety CO39 and barley (*Hordeum vulgare*) cultivar E9, used for infection assays, were grown as previously described.^40^

### 2.2 Growth inhibition assays

Melatonin (73-31-4, Aladdin, China), isoprothiolane, carbendazim (41814-78-2), thiophanate methyl (23564-05-8), and tricyclazole (10605-21-7) from Bide medicine, China, were dissolved in dimethyl sulfoxide (DMSO) (67-68-5, Aladdin, China). The fungicides and their combinations are provided in Tables S1 and S2. Software CompuSyn was used to calculate the combination index.^41–43^

### 2.3 Fungicidal activity assays of melatonin and isoprothiolane

As previously described,^44^ conidial suspensions (2×10^4^ spores/ml) of *M. oryzae* strain P131 supplemented with isoprothiolane, melatonin, or the combination were inoculated on hydrophobic coverslips. The penetration rate was analyzed on barley epidermis.^45^ Conidial germination and appressorium formation were observed and analyzed under the microscope (Nikon Ni90, Japan).^46^

### 2.4 Plant infection assays

Three-week-old rice seedlings and five-day-old barley seedlings were inoculated with conidial suspensions (5×10^4^ spores/ml) in 0.025% Tween-20 of *M. oryzae* strain Guy11 supplemented with DMSO, melatonin, isoprothiolane, or the combination, as previously described.^47^ Symptoms were examined at 7 days post-inoculation (dpi) and 6 dpi for rice and barley, respectively. Drops of conidial suspensions were inoculated onto barley leaves and photographed at 5 dpi. The lesion area was statistically analyzed using ImageJ.^48^

### 2.5 Residual isoprothiolane analysis

The isoprothiolane residues were measured as previously described.^49–51^ Briefly, rice seedlings were sprayed with 17.2 µM isoprothiolane combined with or without 5 mM melatonin, and samples were collected at 1-, 7-, 14-, and 21-dpi. The more detailed information is provided in the Supporting Information.

### 2.6 RNA-seq and reverse transcription quantitative-PCR (qRT-PCR) assays

*M. oryzae* was cultured in liquid CM supplemented with 0.5 % DMSO, 17.2 µM isoprothiolane, 5 mM melatonin, or 5 mM melatonin combined with 17.2 µM isoprothiolane. After 24 h, mycelia were collected for RNA-seq analysis as previously described.^46,52^ The more detailed information is provided in the Supporting Information.

### 2.7 Lipidomics analysis

Mycelia were treated the same as that for transcriptomics assays. After 24 h, the mycelia were collected and freeze-dried. Total lipids extraction methods as previously described.^46^ And lipids were measured with an Agilent HPLC system coupled with a triple quadrupole/ion trap 4000 QTrap mass spectrometer (Applied Biosystems) as previously described.^55^

### 2.8 Model docking

The crystal structure of protein MoIcl1 (PDB: 5e9f) was used. The 3D structures of melatonin and isoprothiolane were retrieved from the PubChem Compound database (https://www.ncbi.nlm.nih.gov/pccompound/). AutoDock Vina was used to dock melatonin and isoprothiolane into protein MoIcl1, and conformations with the best affinity were selected as the final docking conformations.^56^

### 2.9 Generation of the *icl1* mutant of *M. oryzae*

The *Moicl1* (MGG_04895) mutant strain was generated and verified through homologous recombination as described.^57^ Putative knockout mutants were verified by PCR (Table S3).

### 2.10. Protein purification and western blotting

Expression of MoIcl1-6×His was performed as previously described.^56^ The 6×His-tagged protein was purified with Ni NTA Beads 6FF (Smart-Lifesciences Biotechnology, China).^58^ The purified protein was concentrated with 10 kDa ultrafiltration spin columns. For western blotting assays, total proteins were extracted and examined as previously described.^59^ The blotting was performed on the PVDF membrane (Bio-Rad, USA) and imaged with the Western Blotting Detection Kit (Advansta, USA) on the ChemiDoc Touch Imaging System (BIO-RAD, USA).

### 2.11. Surface plasmon resonance (SPR) assays

For SPR assays, the possible melatonin-MoIcl1 interaction was assayed on Biacore T200 (Ge Healthcare Bio Sciences Ab, USA) as previously described.^13^ Briefly, MoIcl1 was immobilized to the CM5 sensor chip surface via amine coupling. Melatonin at concentrations of 62.5-, 125-, 250-, 500-, and 4000 µM was used for SPR assays together with the control.

### 2.12. Differential scanning fluorimetry using intrinsic protein fluorescence (nano DSF)

For nano DSF assays, the possible isoprothiolane-MoIcl1 interaction was assayed on Prometheus NT.48 (NanoTemper, Germany) as previously described.^60^ The more detailed information is provided in the Supporting Information.

### 2.13 Measurement of endogenous melatonin in rice

The endogenous levels of melatonin were measured as previously described.^61^ Briefly, rice seedlings CO39 were sprayed with 5 mM melatonin, 17.2 µM isoprothiolane and 17.2 µM isoprothiolane combined with 5 mM melatonin, and samples were collected at 48 hpi. The more detailed information is provided in the Supporting Information.

### 2.14 Statistical analysis

Data are shown as the mean ± SD. All the measurements were taken from independent samples as indicated. Statistical significance was assessed using the unpaired Student’s *t-*test. Statistical analyses comprising calculation of degrees of freedom were done using GraphPad Prism 6.0. *P* value < 0.05 was considered statistically significant.

## 3 RESULTS

### 3.1 Melatonin and isoprothiolane synergistically inhibited growth of *M. oryzae*

We chose fungicides commonly used in rice blast control to combine with melatonin for possible antifungal synergy (Figure 1A).^62^ In contrast to isoprothiolane or melatonin treatment alone, the inhibitory effects of the combination on hyphal growth were increased (Figure 1B,D). For example, the inhibition rates of hyphal growth by 2 mM melatonin or 1 mg/l isoprothiolane were 24% or 32%, respectively. When combined, the inhibition rate increased to 58%, which is greater than the sum of the individual inhibition rates. In treatments of melatonin combined with carbendazim, thiophanate methyl, or tricyclazole, the inhibitory effects of these combinations were less than the sum of the inhibition rates when used individually (Figure 1C,E,F). Combination index (CI) curve analyses showed that only the CI values of melatonin combined with isoprothiolane were less than 1, indicating the synergy of melatonin and isoprothiolane in inhibiting fungal growth of *M. oryzae* (Figure 1G).

**FIGURE 1.**
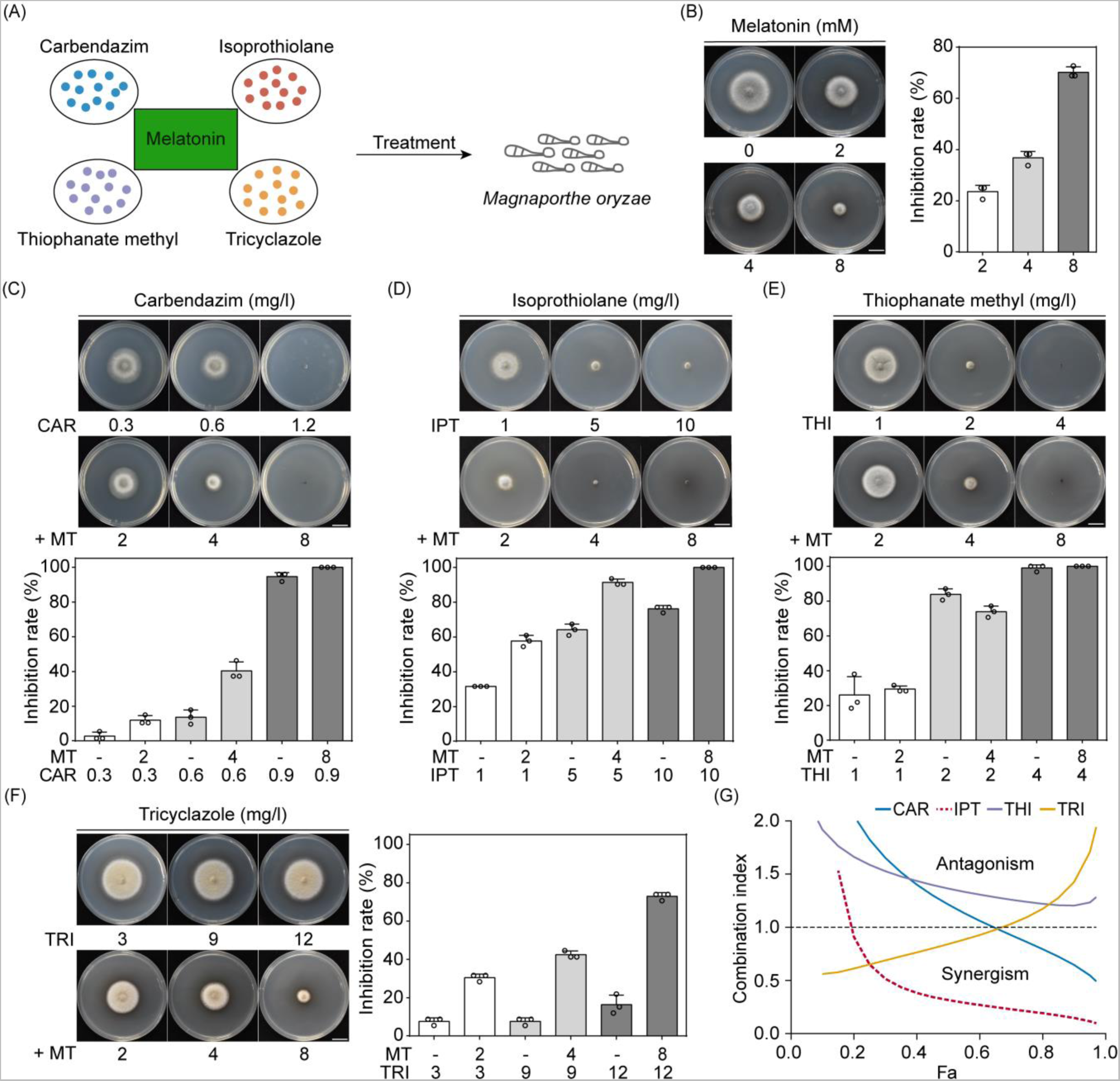
Melatonin (MT) and isoprothiolane (IPT) synergistically inhibit vegetative growth of *Magnaporthe oryzae*. (A) Schematic diagram of the chemical combinations. (B-F) Colony morphology and growth inhibition of *M. oryzae* on complete medium (CM) plates supplemented with MT, different fungicides, and MT combined with fungicides at 5 days post-inoculation (dpi). CAR, carbendazim; THI, thiophanate methyl; TRI, tricyclazole. The inhibition rate (%) = [(diameter of untreated colony − diameter of treated colony)/diameter of untreated colony] × 100. Means and standard deviations were calculated from three independent replicates. (G) Fraction affected (Fa)-combination index (CI) analysis of different MT-fungicide combinations. CI < 1, synergism; CI = 1, additive effect; CI > 1, antagonism. Bars, 1 cm.

### 3.2 Melatonin and isoprothiolane synergistically inhibited development and infection processes of *M. oryzae*

As the concentration increased, the inhibitory effect of melatonin or isoprothiolane on hyphal growth increased (Figure 2A,B,C). In the combination of melatonin and isoprothiolane, the colony diameters were much smaller than that treated with individual chemicals. The inhibitory concentration of 1 mM melatonin combined with 1.7 µM isoprothiolane (combination A) or 3 mM melatonin combined with 1.7 µM isoprothiolane (combination B) were further enhanced compared with that treated with the individual chemicals (Figure S1). Additionally, the CI value of combination A was lower than that of combination B (Figure 2D), suggesting that combination A is the optimal combination among the tests.

**FIGURE 2.**
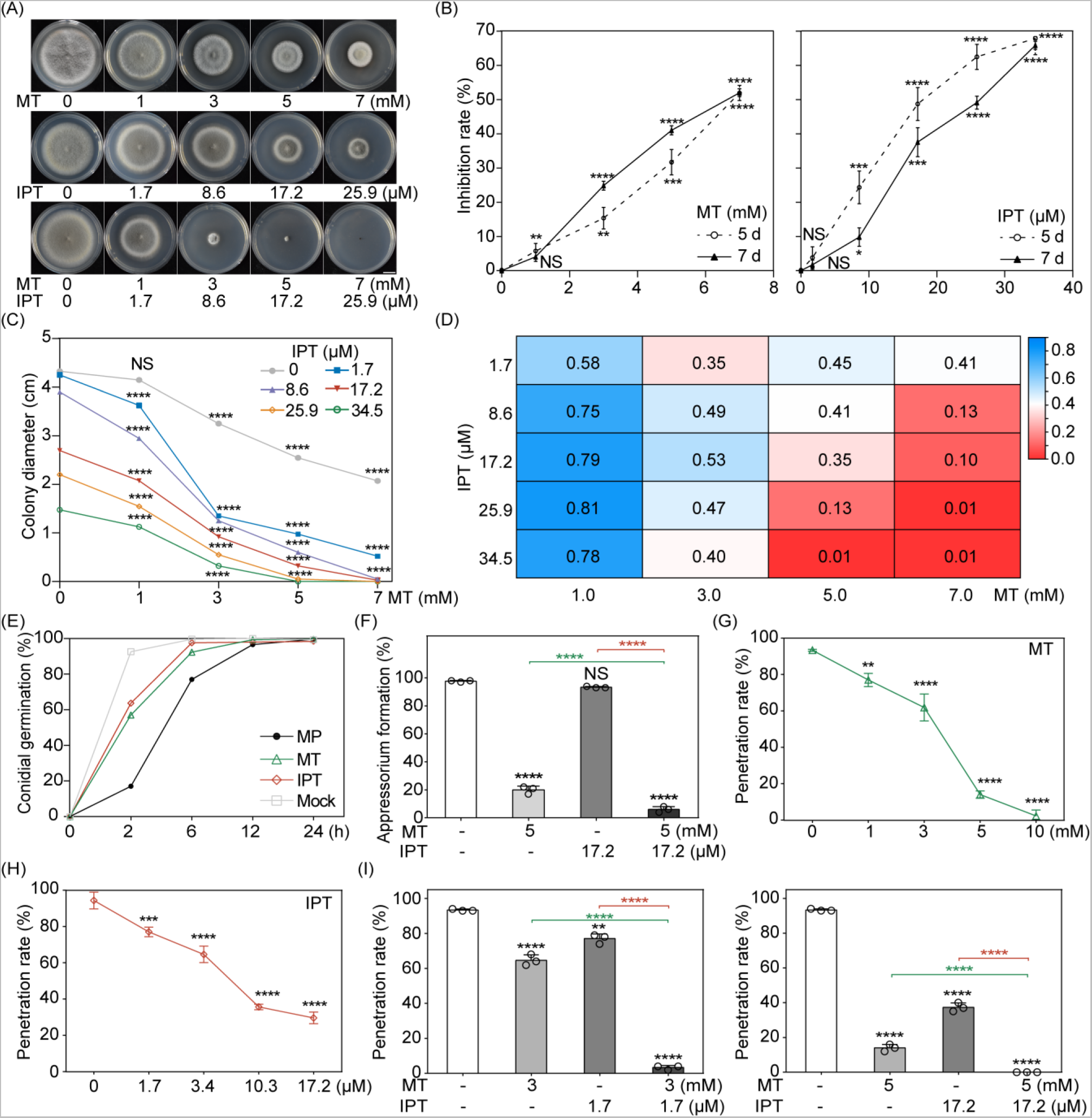
Melatonin (MT) and isoprothiolane (IPT) synergistically inhibit growth and development of *M. oryzae*. (A) Colony morphology of M. oryzae on CM plates supplemented with dimethyl sulfoxide (DMSO), MT, IPT, or melatonin together with isoprothiolane (MP) at 7 dpi. Bar, 1 cm. (B) Fungal growth was inhibited by MT or IPT at 5 and 7 dpi. The inhibition rate (%) = [(diameter of untreated colony – diameter of treated colony)/diameter of untreated colony] × 100. (C) Fungal growth of *M. oryzae* was treated with MP at 7 dpi. (D) The synergistic effect (CI < 1) of MT and IPT. (D) Conidial germination was inhibited by MT, IPT, or MP, respectively at 2 hpi. (E) Appressorium formation was inhibited by MT, IPT, or MP, respectively at 24 hpi. (G-I) Penetration rates on barley by *M. oryzae* treated with MT, IPT, or MP at 36 hours post-inoculation (hpi). Three biological replicates were used for each sample. Error bars represent maximum and minimum values, respectively, and asterisks indicate significant differences using the unpaired Student’s *t*-test. The black asterisk represents comparison with the control, the red asterisk represents a comparison between MP and IPT, and the green asterisk represents a comparison between MP and MT. (\**P* < 0.05, \*\**P* < 0.01, \*\*\**P* < 0.001, \*\*\*\**P* < 0.0001); NS, not significant.

At 2 hours post-inoculation (hpi), the inhibition rates of conidial germination by 1 mM melatonin or 3.4 µM isoprothiolane were 43% or 36%, respectively. The inhibition rate of conidial germination by their combination was high at 83% (Figure 2E), indicating the synergistic inhibitory effect on conidial germination by the combination. Also, this synergistic inhibitory effect was observed in appressorium formation and fungal penetration (Figure 2F-H). When tested individually, the inhibition rates of fungal penetration by 3 mM melatonin and 1.7 µM isoprothiolane were 38% and 23%, respectively, while the combination was at97%, higher than the sum of individual chemicals (Figure 2I). These results together demonstrate that melatonin and isoprothiolane synergistically inhibit infection-related morphogenesis of *M. oryzae*.

### 3.3 Melatonin and isoprothiolane synergistically inhibited plant infection by *M. oryzae*

We performed spray inoculation assays on barley and rice seedlings. With 10 mM melatonin or 345 µM isoprothiolane, only tiny disease lesions were observed on rice (Figure 3A,B). In contrast to those treated with single chemicals, the numbers of lesions on seedlings treated with the combination were significantly reduced (Figure 3C,D). The inhibition rate of 5 mM melatonin combined with 17.2 µM isoprothiolane on plant infection was 76% while that of 172 µM isoprothiolane alone was 69%, indicating that the concentration of isoprothiolane can be reduced by 10-fold to achieve the same or improved efficacy on plant infection when supplemented with melatonin. Similarly, the synergistic effect of melatonin and isoprothiolane on fungal infection was also notable on barley (Figure 3E). These results demonstrate that melatonin and isoprothiolane synergistically inhibit plant infection by *M. oryzae*, reducing the dosage of isoprothiolane necessary for rice blast control.

**FIGURE 3.**
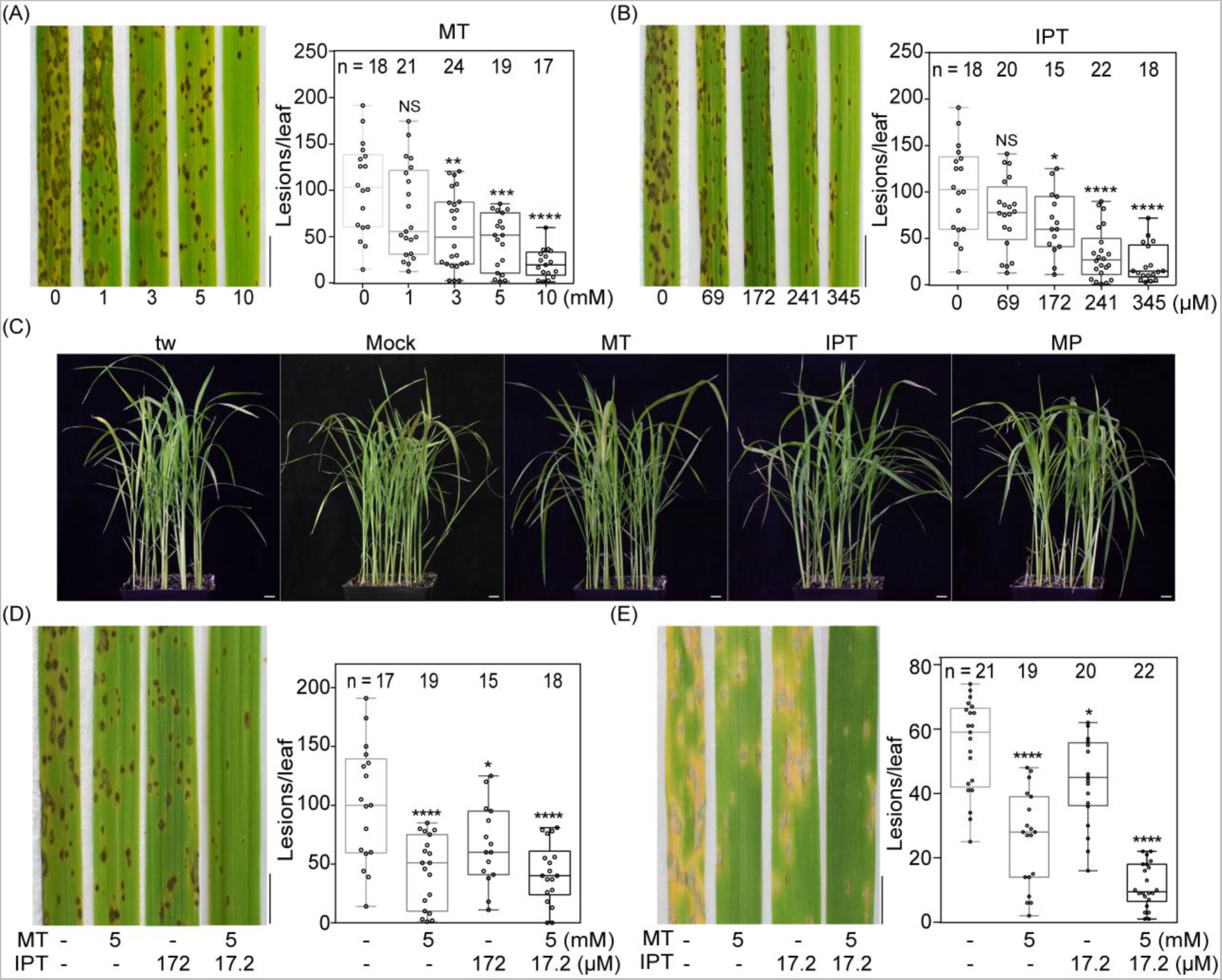
Melatonin (MT) and isoprothiolane (IPT) synergistically inhibit plant infection by *M. oryzae*. (A-B) MT and IPT inhibited fungal infection on rice seedlings at 7 dpi. Bars, 0.5 cm. (C) Spray assays of rice seedlings with conidial suspensions supplemented with DMSO, 5 mM MT, 172 µM IPT, or MP (5 mM MT + 17.2 µM IPT) at 7 dpi. Bars, 1 cm. (D) Representative rice plants and statistical analyses of the lesions from samples shown in (C). Bar, 0.5 cm. (E) Spray assays on barley leaves with conidial suspensions supplemented with DMSO, 5 mM MT, 17.2 µM IPT or MP (5 mM MT + 17.2 µM IPT) at 6 dpi. Bar, 0.5 cm. Error bars represent maximum and minimum values, respectively, and asterisks indicate significant differences using the unpaired Student’s *t*-test (\**P* < 0.05, \*\**P* < 0.01, \*\*\**P* < 0.001, \*\*\*\**P* < 0.0001); NS, not significant.

### 3.4 Melatonin reduced residual levels of isoprothiolane

We used HPLC to measure the residual levels of isoprothiolane in rice leaves treated with or without 5 mM melatonin. In leaves receiving isoprothiolane without melatonin supplementation, 1.819 mg/g isoprothiolane was detectable at 21 dpi. In leaves supplemented with melatonin, the residual level of isoprothiolane was much lower, at 0.344 mg/g, an 81% reduction, indicating that the application of melatonin remarkably reduced isoprothiolane residues in rice (Figure 4).

**FIGURE 4.**
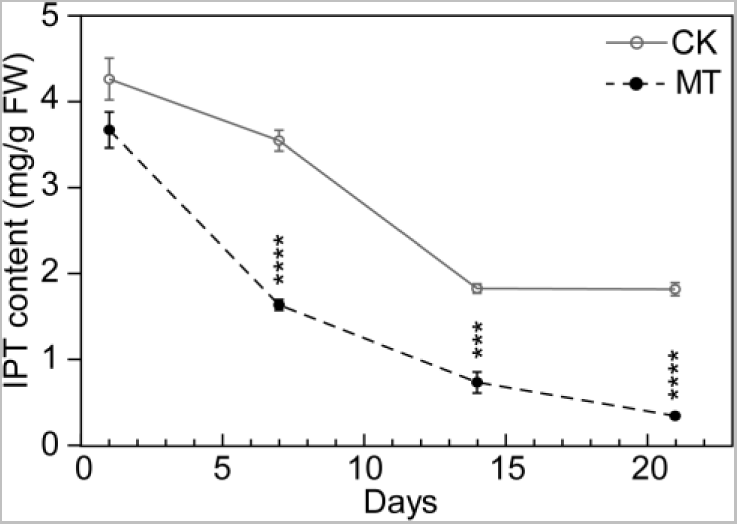
Melatonin (MT) reduces the residual level of isoprothiolane (IPT) in rice. CK, the control treated with isoprothiolane alone; FW, fresh weight. Samples were harvested and analyzed at 1, 7, 14, and 21 days post-application, respectively. Means and standard deviations were calculated from three independent replicates. Data were analyzed with Student’s *t*-test. Asterisks represent significant differences (\*\*\**P* < 0.001, **** *P* < 0.0001).

### 3.5 Melatonin and isoprothiolane increase the endogenous levels of melatonin in rice

We found that the control group was only 0.17 mg/g, but the endogenous levels of melatonin were 0.50 or 0.44 mg/g in rice leaves treated with melatonin or melatonin combined with isoprothiolane (Figure 5B). qRT-PCR assays confirmed that *TDC1*, *ASMT1* and *COMT*, genes involved in melatonin biosynthesis were upregulated, and the change of these genes was greater than that of *M3H*, gene of the melatonin metabolic pathway (Figure 5A,C). These results show that exogenous application of melatonin or melatonin combined with isoprothiolane increases the level of melatonin in rice.

**FIGURE 5.**
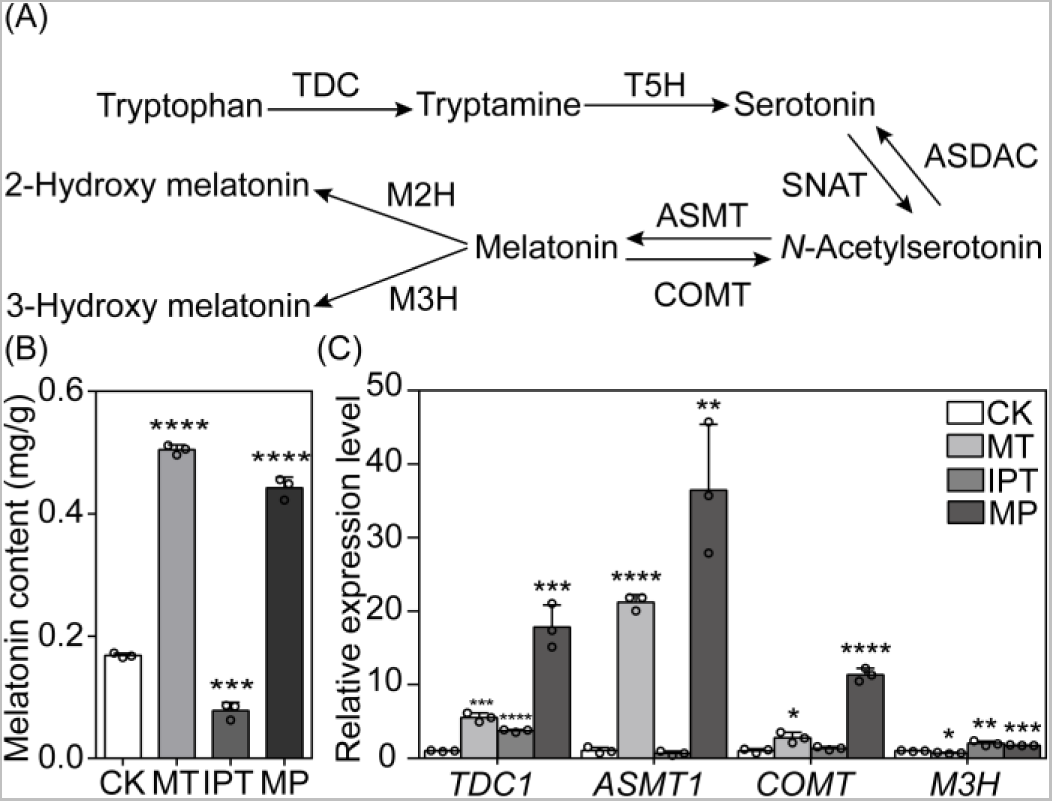
The levels of endogenous melatonin (MT) and genes involved in melatonin biosynthesis in rice. (A) The predicted pathway of melatonin biosynthesis and catabolism in rice. (B) The endogenous levels of melatonin in rice leaves treated with 5 mM MT, 17.2 µM isoprothiolane (IPT) or 5 mM melatonin combined with 17.2 µM isoprothiolane (MP) at 48 hpi. CK, the rice leaves treated with DMSO. (C) qRT-PCR assays of genes showed in (A). Means and standard deviations were calculated from three independent replicates. Data were analyzed with Student’s *t*-test. Asterisks represent significant differences (\**P* < 0.05, \*\**P* < 0.01, \*\*\**P* < 0.001, **** *P* < 0.0001).

### 3.6 Melatonin and isoprothiolane co-suppressed many overlapping targets in *M. oryzae*

We performed RNA-sequence analyses to further investigate the synergistic mechanism of melatonin and isoprothiolane on *M. oryzae*. In total, 1,111 common differentially expressed genes (DEGs) in three mycelial samples treated with melatonin, isoprothiolane, or melatonin combined with isoprothiolane (Figure 6A). These common DEGs were mainly down-regulated, which are enriched in transcriptional process, glucose and lipid metabolism, and oxidation-reduction process (Figure 6B,D). The alterations of gene expression levels of these common DEGs in the group treated with melatonin and isoprothiolane together were greater than those treated with melatonin or isoprothiolane alone (Figure 6C). Among down-regulated DEGs, genes *MoABC3*, *MoCBL4*, *MoCBP1*, *MoC19TRF2*, *MoGEL3*, *MoHEG16*, *MoZFP7*, and *MoZFP14* were reported to regulate infection-related morphogenesis in *M. oryzae*.^63–70^ Consistently, the expression levels of those genes were significantly down-regulated as revealed in qRT-PCR assays (Figure 6E and Figure S2). These results suggest that melatonin and isoprothiolane synergistically inhibit infection-related morphogenesis of *M. oryzae* by co-regulating many genes.

**FIGURE 6.**
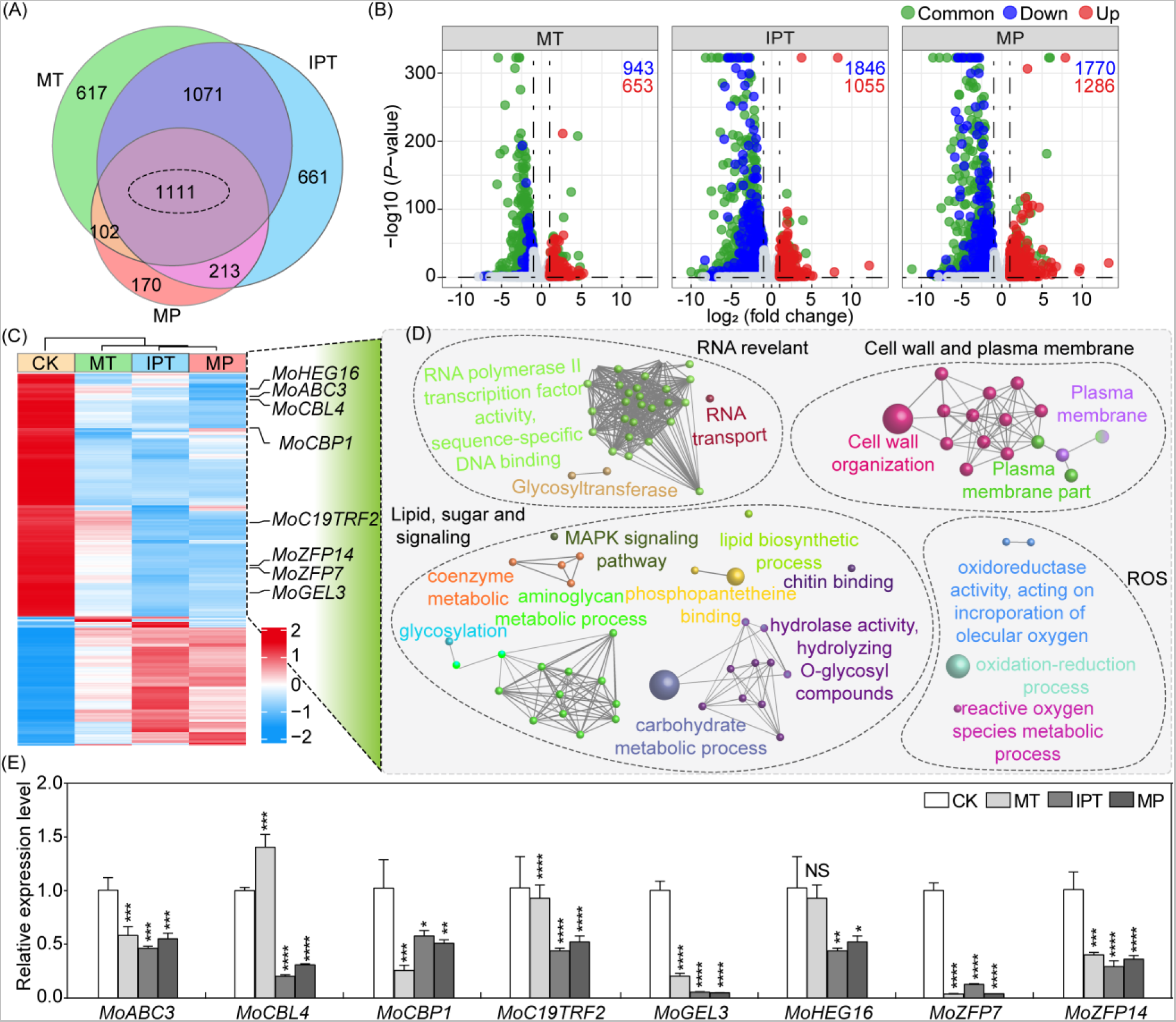
Transcriptomic analysis of the synergy of melatonin (MT) and isoprothiolane (IPT) in *M. oryzae* inhibition. (A) Venn diagram of differentially expressed genes (DEGs) caused by MT, IPT, or melatonin combined with isoprothiolane treatment (MP). (B) Volcano plot of DEGs derived from *M. oryzae* treated with MT, IPT, or MP. (C) Heatmap of common DEGs from samples shown in (b). CK, the control group. (D) GO and Kyoto Encyclopedia of Genes and Genomes (KEGG) enrichment analyses of DEGs visualized with Cytoscape Enrichment Map. GO terms, visualized as colorful dots, with overlapping genes are connected with lines. (E) qRT-PCR assays of DEGs from (C) related to vegetative growth, conidiation, appressorium formation, and pathogenicity. Means and standard deviations were calculated from three independent replicates. Error bars represent maximum and minimum values, respectively, and asterisks indicate significant differences using the student’s *t*-test (\**P* < 0.05, \*\**P* < 0.01, \*\*\**P* < 0.001, \*\*\*\**P* < 0.0001); NS, not significant.

### 3.7 Melatonin and isoprothiolane co-interfere with lipid metabolism in *M. oryzae*

In molecular docking with the crystal structure of MoIcl1 (PDB: 5e9f),^56^ amino acid residues Asp167 and Lys135 of MoIcl1 were predicted to bind to melatonin and isoprothiolane, respectively, through hydrogen bonds with the binding energy at −10.5 kcal/mol (Figure 7A). We expressed and purified MoIcl1 (Fig. S4). SPR assays showed that melatonin binds to MoIcl1, and the dissociation equilibrium constant *K_D_* is 287 µM (Figure 7B,C). Nano DSF showed that thermal transition (unfolding) temperature (Tm) of MoIcl1 protein changed 0.7 ℃ compared with the control, higher than 0.4 ℃, indicating that isoprothiolane enhances the structural stability of MoIcl1 protein (Figure 7D,E).^71^ These results show that MoIcl1 is a common target of melatonin and isoprothiolane.

**FIGURE 7.**
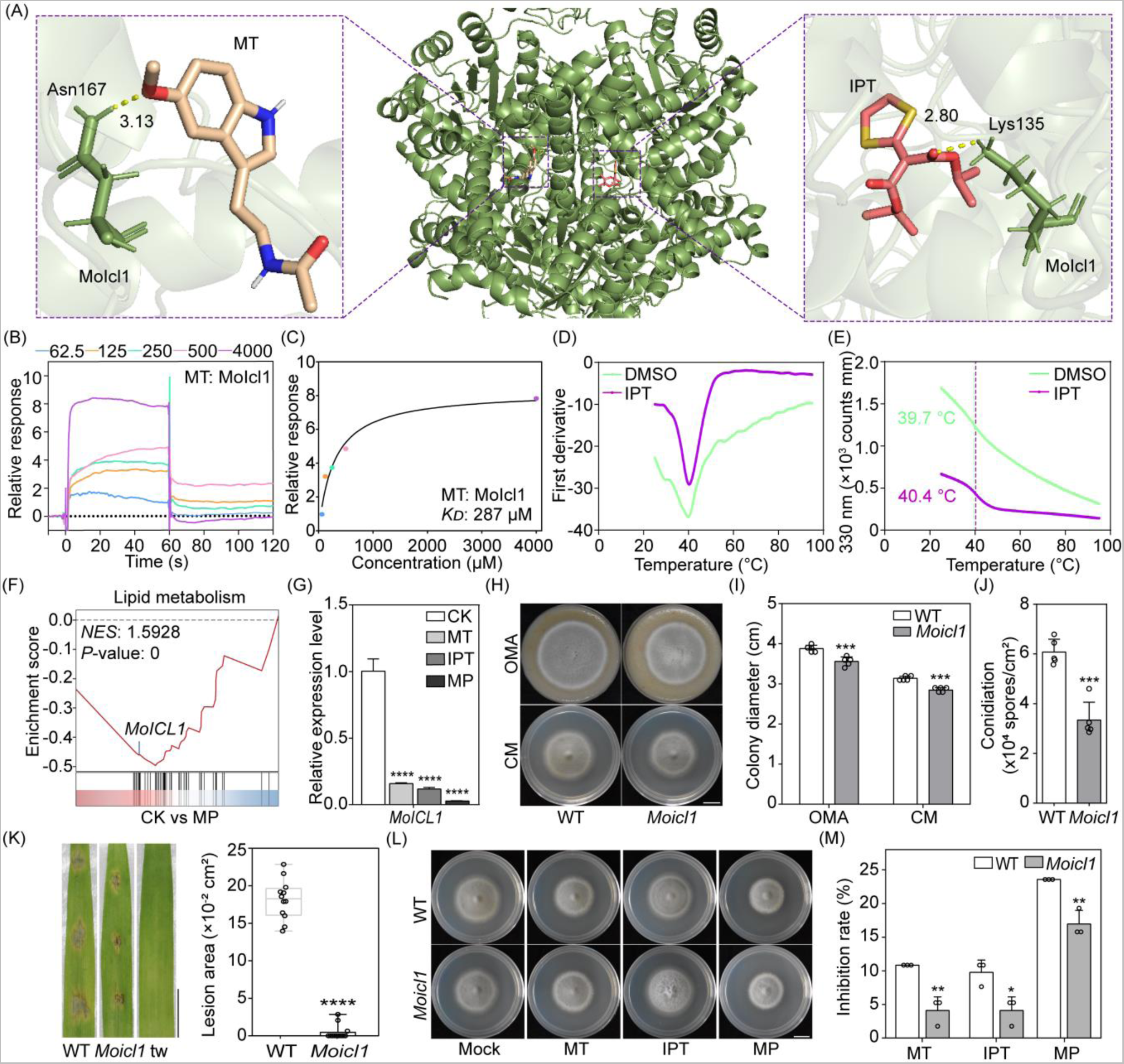
Melatonin (MT) and isoprothiolane (IPT) target protein MoIcl1 in *M. oryzae*. (A) The molecular docking model of MoIcl1 (PDB: 5e9f) with MT and IPT. On the two sides are enlarged views showing the molecular models of MT (milky white) and IPT (brick red) in the binding pocket of protein MoIcl1. (B) Binding of small molecule compound MT to protein MoIcl1. Biacore sensorgrams showing the association and dissociation curves for MT-MoIcl1 binding at indicated concentrations in surface plasma resonance (SPR) assay. (C) The binding affinity *K_D_* was obtained by fitting to steady state model. *K_D_* represents the concentration of MT at half saturation of the SPR signal. (D) The thermal transition (unfolding) temperature (Tm) of MoIcl1 in PBS treated with DMSO (green) or IPT (purple). (E) Differential scanning fluorimetry using intrinsic protein fluorescence (nano DSF) scans for MoIcl1 treated with DMSO (green) or IPT (purple) at 330 nm fluorescence. (F) GSEA enrichment plot of DEGs derived from samples treated with MP associated with lipid metabolism. *NES*, normalized enrichment score. (G) qRT-PCR assays of the *MoICL1* gene. (H) Seven-day-old Guy11 (WT) and *Moicl1* mutant cultures on oatmeal agar (OMA) and complete medium (CM) plates at 28 ℃. Bar, 1 cm. (I) Colony diameters of the strains shown in (H) were calculated from five biological replicates. (J) Conidiation of the WT and *Moicl1* strains from seven-day-old OMA cultures. Means and standard deviations were calculated from five independent replicates. (K) Drop-inoculation assays of WT and *Moicl1* on barley leaves with conidial suspensions (1×10^5^ conidia/ml) at 5 dpi. tw, the control treated with 0.025% Tween-20. Bar, 1 cm. Means and standard deviations were calculated from 12 independent replicates. (L) Seven-day-old cultures of the WT and *Moicl1* mutant strains on CM plates supplemented with 0.5 mM MT, 17.2 µM IPT, or 1 mM melatonin + 1.7 µM isoprothiolane (MP). Inhibition rates are shown in (H). Bar, 1 cm. Means and standard deviations were calculated from three independent replicates. (I) Asterisks indicate significant differences using the student’s *t*-test (\**P* < 0.05, \*\**P* < 0.01, \*\*\*\**P* < 0.0001).

The Gene Set Enrichment Analysis (GSEA) showed that the lipid metabolic pathway was significantly suppressed in the combination treatment (Figure 7F). Lipidomics assays showed that upon treatment with melatonin or isoprothiolane, most lipid biosynthesis processes were inhibited (Figure S3). Taken together, both melatonin and isoprothiolane appear to interfere with lipid metabolism in *M. oryzae*. In particular, the expression of the isocitrate lyase-encoding gene *MoICL1* was significantly reduced under all the above treatments, which was confirmed by qRT-PCR assays (Figure 7G). To analyze the role of *MoICL1* in the combination treatments, we obtained the *Moicl1* mutants.^72^ Deletion of *MoICL1* reduced fungal growth, conidiation, and plant infection by *M. oryzae* (Figure 7H-K). Furthermore, the inhibition rates of melatonin, isoprothiolane, and their combination on mutant *Moicl1* were significantly reduced compared to that of the wild-type (Figure 7L,M). These results show that melatonin and isoprothiolane inhibit the virulence of *M. oryzae* by co-regulating MoIcl1, one of the common targets.

## 4 DISCUSSION

This study aimed to explore an environmentally conscious method to effectively control rice blast, a devastating disease^73,74^ Our results revealed a synergistic inhibition of rice blast by melatonin combined with isoprothiolane. Indeed, the inhibition rates of melatonin combined with isoprothiolane on fungal growth, conidial germination, appressorium formation, and plant infection are higher than the sums of inhibition rates by each chemical. This study is the first to use the combination of melatonin and isoprothiolane to control rice blast and provides evidence of the potential mechanism.

Our study together with others shows that melatonin is an effective antifungal and biofungicidal adjuvant for *M. oryzae* and over 10 other plant pathogens.^9,10,13,75,76^ Plant metabolites such as chitosan, jasmonic acid and pipernonaline also have the inhibition effects on *M. oryzae*.^77–79^ Though environment-friendly, natural small molecules alone often cannot fully satisfy the practical application in the field, the combination of melatonin and isoprothiolane demonstrated here has the potential to solve the above shortcoming of natural molecules in disease control. The synergistic effect of melatonin and isoprothiolane in controlling rice blast are in agreement with observations that drug synergism can improve antifungal efficiency, reduce drug dosage and expand drug targets, possibly leading to delayed drug resistance.^80–83^ In plant infection, adding melatonin to isoprothiolane has the potential to reduce the dosage of isoprothiolane by over 10-fold for rice blast control. Similarly, the dosage of ethylicin can be reduced by 10-fold when supplemented with melatonin in inhibiting vegetative growth of *Phytophthora nicotianae*, the causal agent of oomycetic black shank disease in tobacco.^84^ Multiple other examples have been reported in agriculture and medicine on the synergy of melatonin with various other drugs.^33,35,85^ The possible other fungicides synergistic with melatonin can be predicted.^82^ Melatonin and isoprothiolane may co-suppress related biological processes through overlapping targets, supported by our transcriptomic results that 1,111 DEGs are common in the three treatment groups, and similar synergistic mechanisms have been reported for berberine and eugenol that synergistically depolarize mitochondrial membranes of yeast.^82^

Multiple lines of evidence support our hypothesis that melatonin and isoprothiolane synergistically interfere with lipid metabolism. Many DEGs from samples treated with melatonin together with isoprothiolane are involved in lipid metabolism (Figure 5). Additionally, transcriptomics and lipidomics analyses further support this hypothesis (Figure 7, Figure S3), consistent with reports that melatonin functions through interfering with lipid metabolism in the inhibition of breast cancer.^2,3^ The interactions between MoIcl1 and melatonin as well as isoprothiolane are also confirmed. Furthermore, deletion of one of the common down-regulated DEGs, *MoICL1*, a gene involved in lipid metabolism^87–89^, caused *M. oryzae* less susceptible to melatonin and isoprothiolane. Melatonin and isoprothiolane together resulted in delayed lipid metabolism, reduced growth, conidial germination, appressorium formation, penetration, and plant infection, most of which resemble phenotypes of mutant *Moicl1*.^90^ All these lines of evidence support the conclusion that melatonin and isoprothiolane simultaneously target *MoICL1*, among many DEGs. Protein MoIcl1 is an enzyme not homologous to identified melatonin receptors Cand2, Mt1 or Mt2.^91,92^ In previous studies, we found that Mps1 is one target of melatonin that is important for the appressorium penetration and plant infection, similar to gene *MoICL1*.^13^ Moreover, many other targets are still to be identified as drug synergism often functions through multiple targets.^93^

Interestingly, we found that melatonin reduces the residual levels of isoprothiolane in rice (Figure 4), similar to the observations that melatonin reduces the residual levels of fungicide carbendazim in tomato, which may be caused by regulating redox homeostasis in plants and promoting its metabolism level by enhancing glutathione dependent detoxification.^94^ In this study, the reason for the reduced levels of isoprothiolane may be similar. Under the isoprothiolane treatment, the endogenous level of melatonin is decreased slightly, compared to the control (Figure 5). At the same time, the endogenous levels of melatonin were lower in rice leaves treated with melatonin and isoprothiolane than that treated with melatonin alone due to the negative effects of isoprothiolane on melatonin levels. The possible reason is that melatonin is involved in reducing residues of isoprothiolane in rice and some melatonin is consumed. In this process, expression levels of rice genes involved in melatonin biosynthesis increase to likely synthesize more melatonin to facilitate the detoxification.^94,95^ Additionally, melatonin has been reported to promote plant growth, indicating that the role of melatonin in rice growth is worthy of further studies as one additional potential benefit of melatonin.^96,97^ Melatonin can also improve potassium deficiency tolerance of wheat, showing that melatonin has the potential to improve the resistance of plants to abiotic stress as well.^98^ The efficacy of the fungicide combination may depend on environmental conditions, including temperature, light, humidity, and the timing of applications. Thus, additional studies are needed before large-scale applications are in place. Fortunately, melatonin is commercially available at a low cost at 90-140 USD/kg, which indicates the economic feasibility of the combination. In summary, melatonin combined with isoprothiolane would be a good pair for rice blast control. In future, it is feasible to combine melatonin with other fungicides, herbicides, or insecticides to develop novel and environment-friendly comprehensive pesticides. If needed, the molecular structure of melatonin and isoprothiolane can be further modified to improve their efficacy and to fully utilize their synergy in agriculture. ^13,99,100^

## 5 CONCLUSION

This study has demonstrated the synergistic inhibitory role of melatonin and isoprothiolane in controlling rice blast. Thus, melatonin has the potential to reduce the application dosage of isoprothiolane and reduce isoprothiolane residues in rice. Meanwhile, the combination can increase the level of melatonin in rice, which is beneficial to the growth and development of rice. Melatonin and isoprothiolane synergistically interfere with lipid metabolism and down-regulate transcriptional levels of many genes in *M. oryzae*, including *MoICL1*, to inhibit plant infection (Figure 8). Furthermore, the melatonin-MoIcl1 or isoprothiolane-MoIcl1 interactions were confirmed. Finally, results on co-application of melatonin and isoprothiolane in this study will likely provide valuable information for other possible environment-friendly uses of melatonin in agriculture.

**FIGURE 8.**
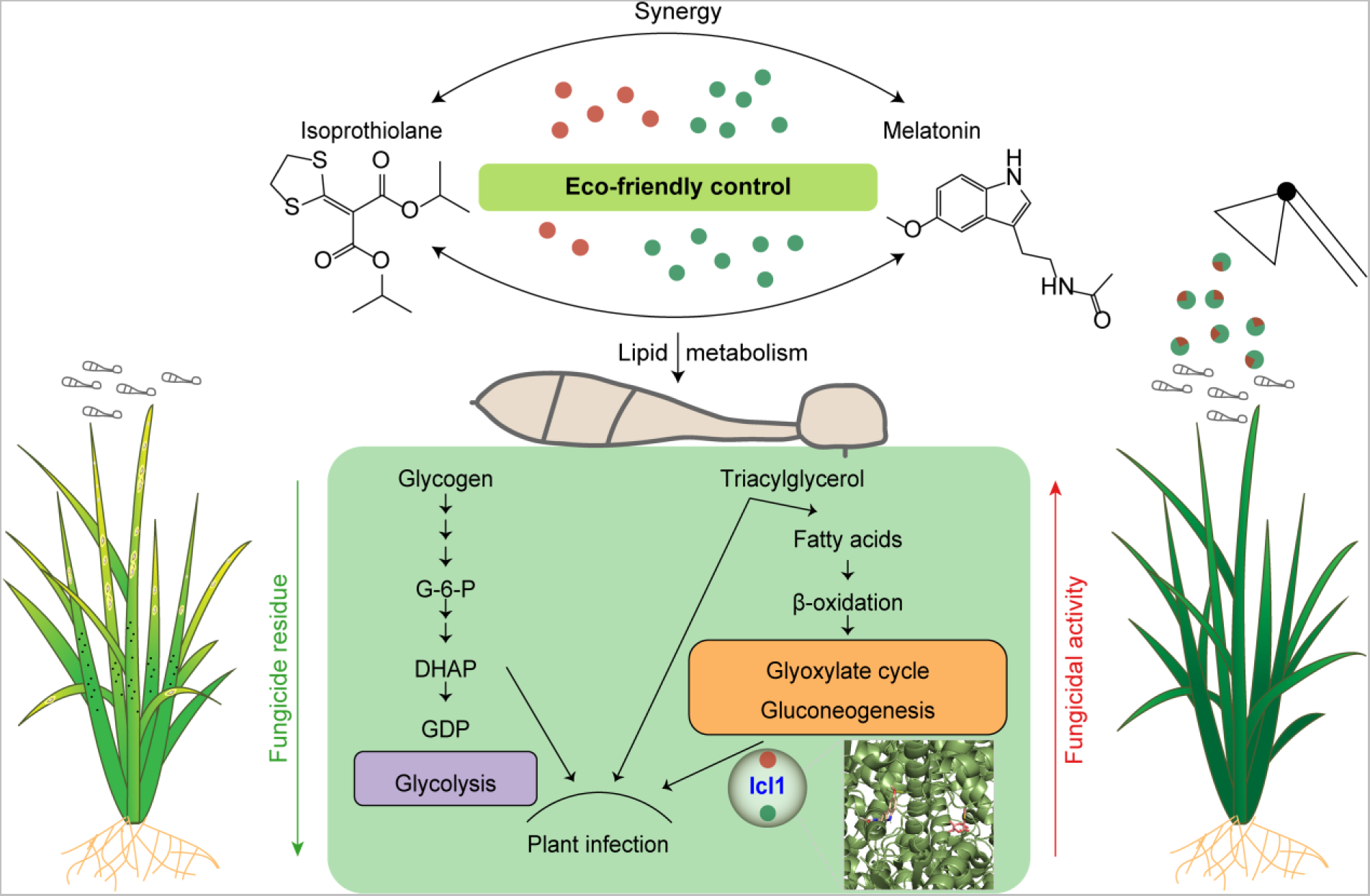
A working model for the synergy of melatonin (MT) and isoprothiolane (IPT) in rice blast control. Melatonin and isoprothiolane interfere with lipid metabolism by regulating multiple targets, including MoIcl1. Meanwhile, the combination of melatonin and isoprothiolane improves fungicidal activity and reduces the residual level of isoprothiolane.

## AUTHOR CONTRIBUTIONS

G.L., R.B. and R.L. designed the experiments. R.B., Z.X., H.C., and W.Y. performed the experiments. G.L., R.L., and R.B. analyzed the data. G.L. and R.B. drafted the manuscript, and G.L., R.B., R.L., Z.X., H.C., J.Z., Y.Z., B.W., P.S., W.Y., L.Z., X-L.C., C-X.L., H.T., and Q.L. revised the manuscript. All authors read and approved the final manuscript.

## ACKNOWLEDGEMENTS

We thank Dr. Larry D. Dunkle at Purdue University for critical reading of this manuscript and Ms. Dongqin Li from the National Key Laboratory of Crop Genetic Improvement for technical assistance in lipidomics analyses. SPR and nanoDSF data were acquired at the State Key Laboratory of Agricultural Microbiology Core Facility. We also thank Gan Sha, Qiping Sun, and other Li lab members for their comments and discussions. This work was supported by National Key Research and Development Program of China (2022YFA1304402), the National Natural Science Foundation of China (31801723, 32172373), and Fundamental Research Funds for the Central Universities ((2023ZKPY002) to G. L. This work was also supported by Hubei Hongshan Laboratory.

## DATA AVAILABILITY STATEMENT

RNA-seq data used in this study can be accessed at the Genome Sequence Archive (GSA) under the accession number CRA008702 (https://ngdc.cncb.ac.cn/gsa). RNA-seq data used in this study can be accessed under the accession number of PRJNA910920 at the National Center for Biotechnology Information (NCBI) (http://www.ncbi.nlm.nih.gov/sra).

## Figure Legends

**FIGURE S1** The inhibitory efficiency on growth of *M. oryzae* by melatonin (MT) combined with isoprothiolane (IPT) compared with individual chemicals. Means and standard deviations were calculated from four independent replicates.

**FIGURE S2** Classification of differentially expressed genes (DEGs) common in samples treated with melatonin, isoprothiolane or melatonin together with isoprothiolane. The four colored circles represent DEGs involved in fungal growth, conidiation, appressorium formation, and pathogenicity, respectively. The number indicates DEGs with one or more functions.

**FIGURE S3** High-performance liquid chromatography/mass spectrometry (HPLC-MS) analysis of lipids in *M. oryzae* treated with melatonin (MT) and isoprothiolane (IPT). CK, the control treated with DMSO. DAG, diacylglycerol; PA, phosphatidic acid; PC, phosphatidylcholine; PE, phosphatidylethanolamine; PG, phosphatidylglycerol; PI, phosphatidylinositol; PS, phosphatidylserine; TAG, triacylglycerol. Means and standard deviations were calculated from three independent replicates. Data were analyzed with Student’s *t*-test. Asterisks represent significant differences (\*\*\**P* < 0.001, **** *P* < 0.0001); NS, not significant.

**FIGURE S4** Expression of MoIcl1-6×His protein in *E. coil* and affinity purified. Total proteins further analyzed by SDS-PAGE and Western blot. The grey triangles indicate the position of MoIcl-6×His band. IPTG indicates the protein expression of MoIcl1 induced by 0.1 mM IPTG. AP indicates affinity purified proteins.

